# GenomeMUSter mouse genetic variation service enables multi-trait, multi-population data integration and analyses

**DOI:** 10.1101/2023.08.08.552506

**Authors:** Robyn L. Ball, Molly A. Bogue, Hongping Liang, Anuj Srivastava, David G. Ashbrook, Anna Lamoureux, Matthew W. Gerring, Alexander S. Hatoum, Matthew Kim, Hao He, Jake Emerson, Alexander K. Berger, David O. Walton, Keith Sheppard, Baha El Kassaby, Francisco Castellanos, Govind Kunde-Ramamoorthy, Lu Lu, John Bluis, Sejal Desai, Beth A. Sundberg, Gary Peltz, Zhuoqing Fang, Gary A. Churchill, Robert W. Williams, Arpana Agrawal, Carol J. Bult, Vivek M. Philip, Elissa J. Chesler

**Author notes:** Corresponding Author: The Jackson Laboratory 600 Main Street, Bar Harbor, ME USA.

## Abstract

Hundreds of inbred laboratory mouse strains and intercross populations have been used to functionalize genetic variants that contribute to disease. Thousands of disease relevant traits have been characterized in mice and made publicly available. New strains and populations including the Collaborative Cross, expanded BXD and inbred wild-derived strains add to set of complex disease mouse models, genetic mapping resources and sensitized backgrounds against which to evaluate engineered mutations. The genome sequences of many inbred strains, along with dense genotypes from others could allow integrated analysis of trait – variant associations across populations, but these analyses are not feasible due to the sparsity of genotypes available. Moreover, the data are not readily interoperable with other resources. To address these limitations, we created a uniformly dense data resource by harmonizing multiple variant datasets. Missing genotypes were imputed using the Viterbi algorithm with a data-driven technique that incorporates local phylogenetic information, an approach that is extensible to other model organism species. The result is a web– and programmatically-accessible data service called GenomeMUSter (https://muster.jax.org), comprising allelic data covering 657 strains at 106.8M segregating sites. Interoperation with phenotype databases, analytic tools and other resources enable a wealth of applications including multi-trait, multi-population meta-analysis. We demonstrate this in a cross-species comparison of the meta-analysis of Type 2 Diabetes and of substance use disorders, resulting in the more specific characterization of the role of human variant effects in light of mouse phenotype data. Other applications include refinement of mapped loci and prioritization of strain backgrounds for disease modeling to further unlock extant mouse diversity for genetic and genomic studies in health and disease.

## INTRODUCTION

Hundreds of inbred mouse strains (NCBI:txid10090) and thousands of intercross progeny have been characterized for disease-related traits, developmental characteristics, age-related phenotypes, microbiota, response to infections and other environmental factors, as well as molecularly characterized endpoints, such as transcriptomes, proteomes and metabolomes performed across many tissues. Advances in genetics, including transcriptome-wide and phenome-wide association analysis methods (Gamazon et al. 2015, Dai et al. 2023, Bastarache et al. 2022) and new methods for integration of data obtained from multiple species (Reynolds et al. 2021) create compelling new opportunities for using fully-reproducible and widely studied inbred mouse strains to characterize the polygenetic basis for individual differences in disease-related traits and to more precisely associate phenotypic variation with genetic variation. These applications require uniformly dense information on genotypic variation across populations.

However, the genomes of most inbred strains and intercross progeny have been very sparsely and non-uniformly characterized through studies performed using a variety of genotyping arrays over the past few decades. In contrast, extensive sequencing efforts for a small number of the most widely used inbred strains have generated a tremendous amount of information about polymorphisms, structural variants, and other types of genetic diversity. These data have been presented in various interfaces to support evaluation of the state of region-specific, known variants across inbred strains (Blake et al. 2021, Mulligan et al. 2017, Keene et al. 2011), but often cannot be accessed programmatically to support genome-wide calculation within or across populations. The problems associated with sparse data, the technical difficulty of merging disparate datasets, the scalability and interoperability of the resources and other challenges thus make it difficult to use these data collectively to perform robust genetic analyses, such as Genome-Wide Association Studies (GWAS), with adequate power and density. Therefore, an integrated data resource is needed.

Prior efforts to harmonize variant data sets and address the sparsity of data incorporated known or inferred similarity of sequence across well-characterized strains into genotype imputation algorithms (Szatkiewicz et al. 2008, Kirby et al. 2010, and Wang et al. 2012). Most prior studies included a small number of strains, represented a relatively narrow subset of mouse diversity, and evaluated sparse genotypes. Furthermore, the algorithms these studies employed require computationally expensive parameter estimation methods and often, rely heavily on close or known ancestral relations among the strains. Szatkiewicz et al. and Kirby et al. trained Hidden Markov Models (HMMs) with various numbers of hidden states, estimating the HMM parameters with expectation-maximization (EM) algorithms (Churchill 1989, Frazer et al. 2007). Szatkiewicz et al. imputed genotypes for 51 strains using the trained HMM and the Viterbi algorithm (Viterbi 1967), which converges on the most probable path, given the observed data. Kirby et al. imputed genotypes with the trained HMM and an EM algorithm, EMINEM (Kang et al. 2010), assembling known and imputed genotypes for 94 strains at 8.27+ million locations. Wang et al. 2012 incorporated phylogenetic information, imputing genotypes for 100 classical inbred strains by using the four-gamete rule (Hudson and Kaplan 1985) to define haplotype blocks and infer locally perfect phylogeny. This requirement for close phylogeny and the need to estimate large numbers of free parameters for the HMMs, an O(n) computation which grows linearly with the number of SNPs (Szatkiewicz et al. 2008), limits the application of these algorithms and makes them unsuitable to a comprehensive merged variant set with sparse genotypes encompassing the extensive diversity of extant mouse strains and their intercross derivatives. Data from prior efforts are also somewhat limited in their analytic applications and presentation in authoritative data resources because the methods employed do not provide a locally precise estimate of the accuracy of imputed results, as would be needed for calculations involving weighted variant states.

Therefore, we present an approach and implemented data service to provide a broad and more comprehensive mouse variant resource. The approach is not limited to classical inbred strains, can be applied in a computationally efficient manner without the need to estimate large numbers of free parameters, incorporates local phylogenetic information over larger regions without prior knowledge of strain relatedness, is sensitive to co-occurring missingness while leveraging aggregate genetic similarity measures to predict the most likely allelic state, and provides a local accuracy estimate that can be used in downstream analyses. We evaluated the imputation accuracy on a ‘hold out’ test set that was not used in the imputation process. We deliver these data in a harmonized, scalable, and programmatically accessible annotated variant resource, GenomeMUSter, which will remain essential even when the majority of strains are sequenced. We present its application to multi-population and multi-species analyses of complex trait variation in Type 2 Diabetes and substance use disorders and compare these results to human genetics studies.

## RESULTS

GenomeMUSter brings together the known sequence variation among *Mus musculus* strains. The data resource includes merged and harmonized allelic state data from sixteen datasets that include two Sanger releases (REL2004 and REL1505; Keene et al. 2011), whole genome sequencing of the Collaborative Cross (CC) (Srivastava et al. 2017), BXD recombinant inbred strains (Sasani et al. 2022, Ashbrook et al. 2022), C57BL/6J Eve (B6Eve) (Sarsani et al. 2019), long read whole genome sequencing of 42 inbred strains (Arslan et al. 2023), and ten SNP genotype datasets previously curated on the Mouse Phenome Database (Kirby et al. 2010, Yang et al. 2011, Frazer et al. 2011, Morgan et al. 2015, Srivastava et al. 2017) (**Table 1**). After merging and harmonizing these 16 variant datasets, GenomeMUSter contains segregating allelic data at 106,811,517 genomic locations across 657 inbred strains and includes genotype data for chromosomes Y and MT across 654 and 483 strains, respectively. Though some of the variant sets have no coverage across chromosomes Y and MT, many of the strain sets overlap among the 16 datasets, providing a more complete genome for most strains (**Fig. 1**) than any single dataset alone. Comparison of the merged and harmonized variant dataset revealed that the largest whole genome sequencing dataset (Sanger REL2004), which is comprised of allelic data for 53 strains at 79.8M sites, accounts for only 74.8% of the 106.8M sites and less than 10% of the 657 strains in the merged dataset, while genotyped datasets contained allelic states for < 1% of the sites in the merged dataset (**Fig. 1**, **Table 1**). The relative density of coverage across variant sets varies greatly overall (**Supplemental Table S1**) and by chromosome (**Supplemental Table S2**).

**Fig. 1.**
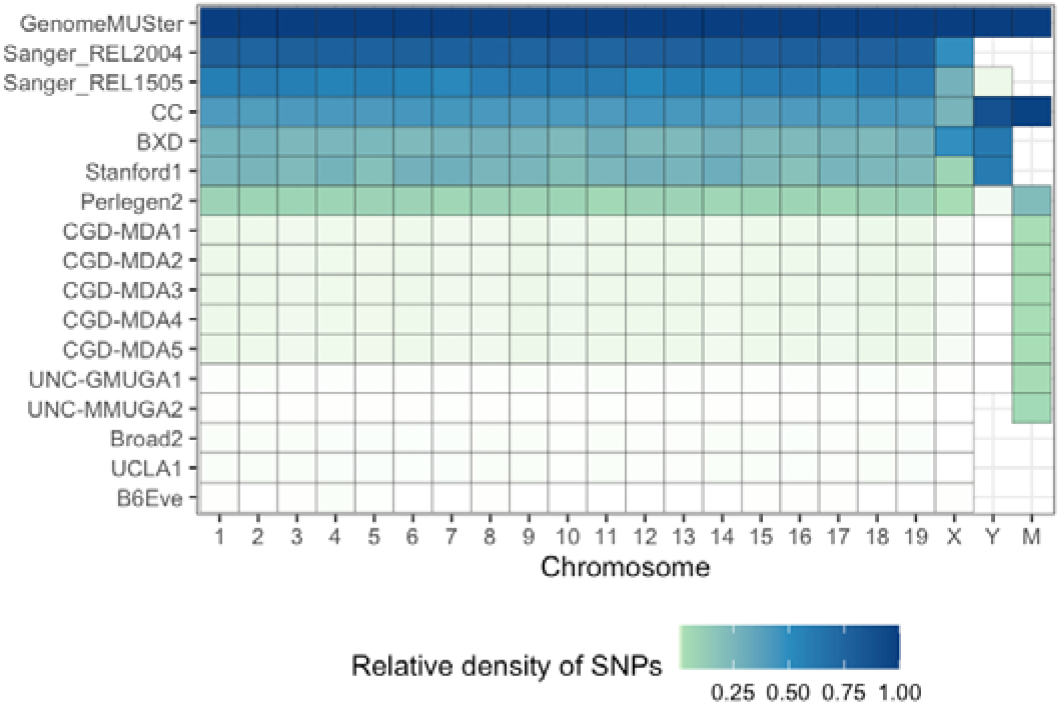
Relative density of information in each dataset to GenomeMUSter. The intensity of the color represents the density of SNPs for each dataset and chromosome, with white, light green, and dark blue representing a density near zero, less than 0.25, and 1, respectively.

**Table 1.** Variant datasets in GenomeMUSter.

| Dataset | Locations | Panel | References |
| --- | --- | --- | --- |
| B6Eve<sup>a</sup> | 56.9K, Chr 1-19, X | Inbred (C57BL/6J Eve) | Sarsani et al. 2019 |
| Broad2 | 131K, Chr 1-19, X | Inbred (89) | Kirby et al. 2010 |
| BXD<sup>a</sup> | 28.6M, Chr 1-19, X,Y | BXD w/parents (151) | Sasani et al., 2022 & Ashbrook et al., 2022 |
| CC<sup>a</sup> | 43.9M, Chr 1-19, X, Y, MT | Collaborative Cross (69) | Srivastava et al. 2017<sup>b</sup> |
| CGD-MDA1 | 470K, Chr 1-19, X, Y, MT | Inbred (142) | Yang et al. 2011 |
| CGD-MDA2 | 470K, Chr 1-19, X, Y, MT | BXD w/ parents (92) | Yang et al. 2011 |
| CGD-MDA3 | 470K, Chr 1-19, X, Y, MT | ILSXISS w/ parents (69) | Yang et al. 2011 |
| CGD-MDA4 | 470K, Chr 1-19, X, Y, MT | AXB, BXA, BXH, CXB, AKXL w/parents (72) | Yang et al. 2011 |
| CGD-MDA5 | 470K, Chr 1-19, X, Y, MT | B6.A, B6.PWD (53) | Yang et al. 2011 |
| Perlegen2 | 8.1M, Chr 1-19, X, Y, MT | Inbred (16) | Frazer et al. 2007 |
| Sanger REL2004<sup>a,c</sup> | 79.8M, Chr 1-19, X | Inbred (53) | Keene et al. 2011<sup>c</sup> |
| Sanger REL1505<sup>a,d</sup> | 59.2M, Chr 1-19, X, Y | Inbred (49) | Keene et al. 2011<sup>d</sup> |
| Stanford1<sup>a</sup> | 25.5M, Chr 1-19, X,Y,MT | Inbred (42) | Arslan et al. 2023 |
| UCLA1 | 132K, Chr 1-19, X | Hybrid Mouse Diversity Panel (248) | e |
| UNC-GMUGA1 | 131K, Chr 1-19, X, Y, MT | Collaborative Cross w/ parents (77) | Morgan et al. 2015 |
| UNC-MMUGA2 | 76K, Chr 1-19, X, MT | Collaborative Cross w/ parents (77) | Srivastava et al. 2017 |

<sup>a</sup>Not an MPD SNP legacy dataset, <sup>b</sup>https://csbio.unc.edu/CCstatus/CCGenomes/, <sup>c</sup>sanger\_38\_rel\_2004\_v7\_dbsnp142,
<sup>d</sup>sanger\_38\_rel\_1505\_v5\_dbsnp142, <sup>e</sup>http://mouse.cs.ucla.edu/mousehapmap/emma.html

Prior to imputation, 90% of strains were missing allelic state information at more than 74% of the sites in the merged dataset (**Fig. 1**), revealing the need for an accurate imputation algorithm. Within chromosome and within each 10Mb region, missing genotypes for each strain were imputed with a data-driven approach that incorporates local phylogenetic information and the Viterbi algorithm, as implemented in HaploQA (https://haploqa.jax.org; **Fig. 2**). HaploQA was designed to perform quality assurance analyses on genotype array data to resolve haplotypes. Here, we take advantage of the computationally efficient implementation of the Viterbi algorithm in the HaploQA software platform along with local phylogenetic information from the merged and harmonized variant dataset to impute genotypes. After imputation, every strain had a known or imputed genotype for at least 84,606,399 sites (> 80% complete allele calls) with median of complete allele calls at 97,442,189 sites (91.6% complete; **Supplemental Tables S3, S4**).

**Fig 2.**
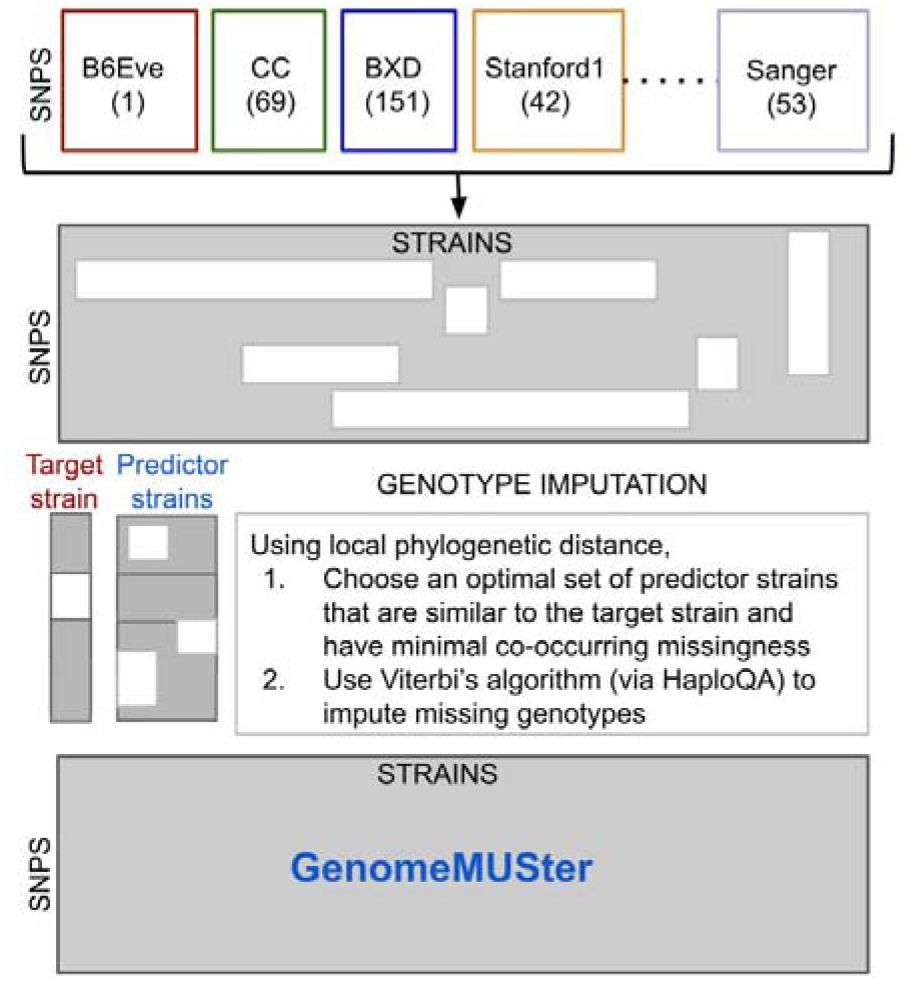
Illustration of the analytical methods used to construct GenomeMUSter. White rectangles represent missing data while gray represents known data. First, sixteen variant datasets were merged and harmonized then missing genotypes were imputed using a data driven approach.

To evaluate the accuracy of the imputation approach for each strain, a “hold out” test set for each strain was randomly selected in each 10Mb region. This test set was not used in the imputation process. Based on aggregated held-out test sets in each 10 Mb region, the median genotype imputation accuracy on the test set across 657 strains was 0.944 with interquartile range [0.923, 0.991] and achieved accuracies greater than 0.80 for 636 of 657 (96.8%) of inbred strains (**Supplemental Tables S3, S4**). The median accuracy across 30 wild-derived inbred strains was 0.769 with interquartile range [0.629, 0.845] yet, not surprisingly, sequence imputation remained low for a few wild-derived inbred strains such as PAHARI/EiJ, CAROLI/EiJ, and SPRET/EiJ, with estimated accuracies and 95% confidence intervals of 0.331 (0.326, 336), 0.485 (0.480, 0.490), and 0.532 (0.529, 0.535), respectively (**Fig. 3, Supplemental Table S3**). These strains, of increasing interest to mammalian geneticists (Chang et al. 2017, Poltorak et al. 2018), represent different *Mus* species. They are distinct from the most deeply characterized *Mus musculus domesticus* strains, which include classical inbred strains, many of which have been sequenced, yielding higher accuracy across the genome (**Fig. 3**).

**Fig 3.**
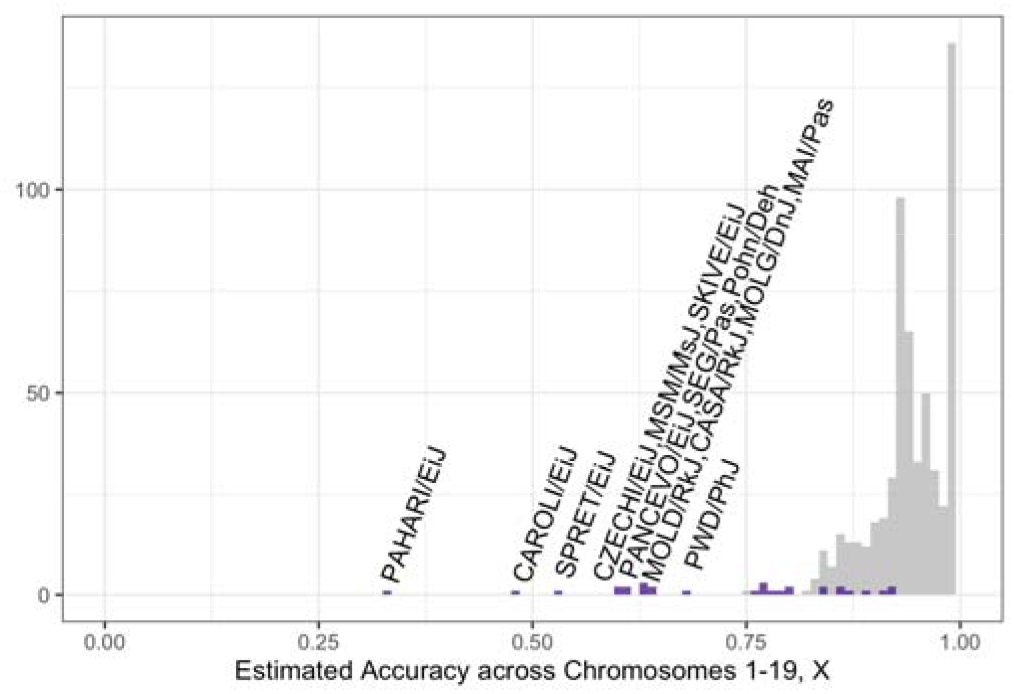
Histogram of estimated imputation accuracies across chromosomes 1-19, and X for 657 strains. Wild-derived strain accuracies are colored purple. Strains with accuracies below 0.70 are annotated.

### GenomeMUSter enables multi-population, multi-trait meta-analysis

A major utility of GenomeMUSter is that it enables genetic discovery across the wealth of recent and historical studies of mouse population variation in an incredibly deep and infinitely extensible collection of complex disease relevant characteristics. We evaluated this capability in the realm of diabetes and obesity research by analyzing a heterogenous set of 276 phenotypic endpoints obtained from 7 studies in the Mouse Phenome Database that measured the response to a high-fat diet, and another set of 122 phenotypic endpoints containing the term ‘glucose’ in their meta content. Studies measured response to a high-fat diet through various methods that include histopathology, lipid profiles and metabolic panels, hormone and metabolite quantification, bone density, and organ and body weights and dimensions. Studies that measured phenotypic endpoints related to glucose used glucose and insulin tolerance methods, metabolic and metabolite panels, and hormone quantification methods. These cross-population analyses on the response to a high-fat diet and endpoints related to glucose were computed on ∼500 Diversity Outbred (J:DO) mice, C57BL/6 consomic strains, Collaborative Cross, Collaborative Cross Pre, the Hybrid Mouse Diversity Panel, and other classical and wild-derived inbred strains (Paigen et al., Svenson et al. 2007, Li et al. 2008, Shockley et al. 2009, Tsuchiya et al. 2012, Morgan et al. 2014, Keller et al. 2018, Lake, et al., Jackson Laboratory, Champy et al. 2008, Center for Genome Dynamics, Lin et al. 2005, Philip et al. 2011, Ghazalpour et al. 2014, Kollmus et al. 2020, Bachmann et al. 2022). Studies in these analyses that measured inbred strains included 6 to 44 strains (median of 14 strains) in the high-fat diet response analysis and 2 to 190 (median of 8 strains) on the analysis across endpoints related to glucose.

We performed a GWAS meta-analysis across these heterogeneous set of measurements to detect variants that affect the high-fat diet response (**Fig. 4A**) and glucose related endpoints (**Fig. 4B**). We identified 1169 variants with effects on high-fat diet response and 1284 variants with effects on glucose related endpoints across this heterogeneous set of measurements; 597 variants with effects in the 99.5^th^ percentile (top peaks) for each analysis and, to account for linkage, an additional 572 variants (high-fat diet response) and 687 variants (glucose) highly correlated with those variants with effects in the 99.5^th^ percentile. These variants are associated with 224 (high-fat diet response) and 287 (glucose) mouse genes and their human orthologs through evidence from PLAC-seq, ChIA-PET, EPD/CAGE, and eQTL results from Diversity Outbred mouse populations (Reynolds et al. 2021). Of the human ortholog genes associated with a high-fat diet response, 93 were associated with 284 Disease Ontology terms (**Supplemental Table S5**). Of the human ortholog genes associated with glucose related endpoints, 128 were associated with 384 Disease Ontology terms (**Supplemental Table S6**). When we evaluated the Disease Ontology annotations of human orthologs of mouse genes associated to the variant effects of exposure to a high-fat diet to genes annotated to human diseases and conditions, Alzheimer’s Disease was the most frequent annotation to the mouse-derived orthologue set, followed by end stage renal disease, asthma, hepatocellular carcinoma, hypertension and obesity and other inflammatory diseases that include Type 2 Diabetes (**Fig. 4C, Supplemental Table S5**). Disease Ontology annotations of human orthologs of mouse genes associated to the variant effects of glucose related endpoints to genes annotated to human diseases and conditions, Type 2 Diabetes was the most frequent annotation to the mouse-derived orthologue set, followed by Alzheimer’s Disease, obesity, and other inflammatory diseases (**Fig. 4D, Supplemental Table S6**). Examining these results together, we find that 148 variants have effects on both the high-fat diet response and glucose related endpoints and these are associated with 43 human ortholog genes. These results indicate that mouse multi-trait meta-analysis produces disease relevant information and further, by examining the precise traits and genes, can facilitate characterization of their role in disease.

**Fig. 4.**
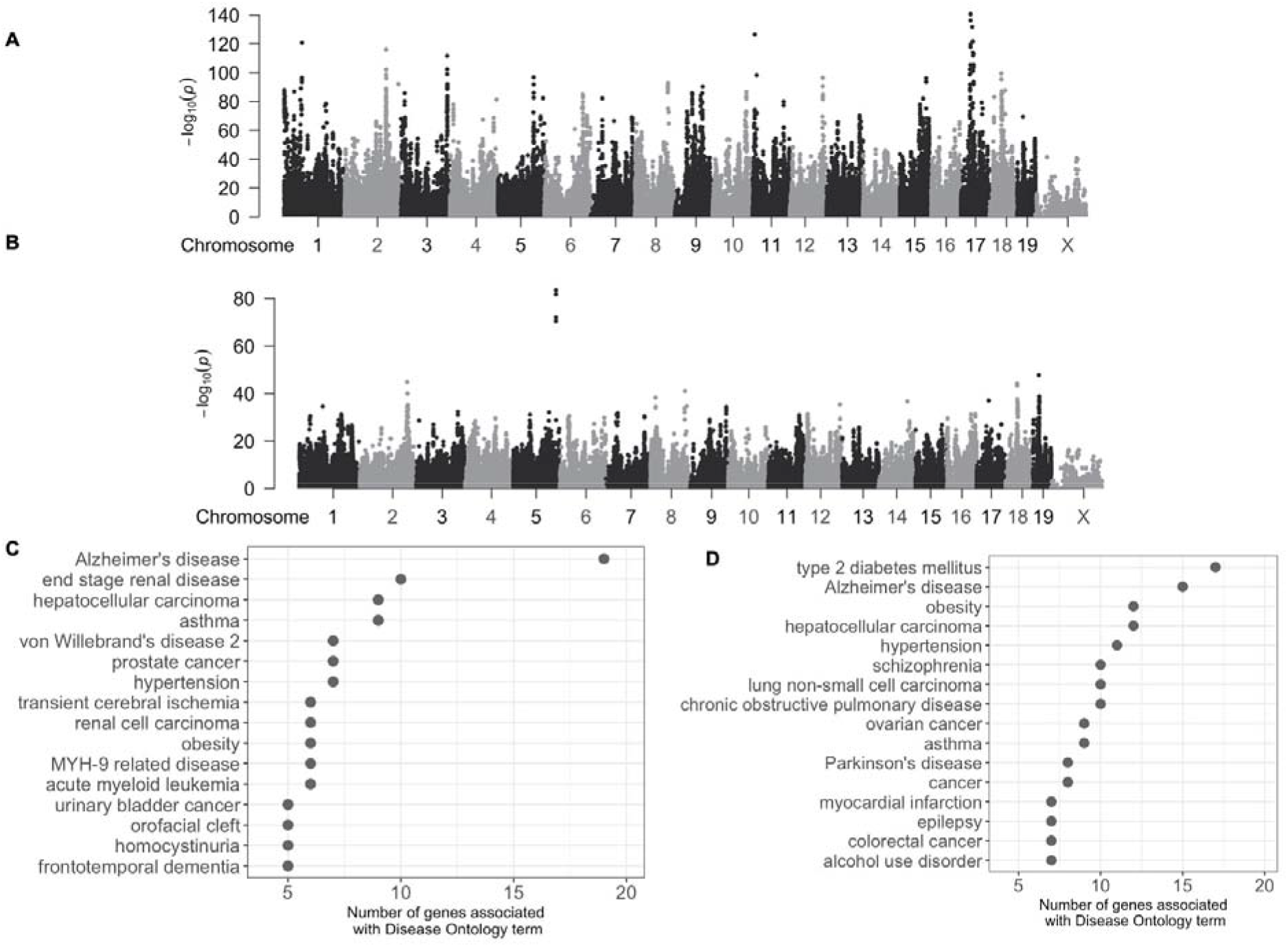
Mouse cross population multi-trait meta-analysis of physiological measurements related to the response to high-fat diet (**A**) and glucose related endpoints (**B**) identify gene orthologs annotated to Alzheimer’s Disease (**C**), Type 2 Diabetes (**D**), and other inflammatory diseases.

### Characterizing Human GWAS with mouse genetic traits

We applied this multi-species approach in the opposite direction, going from human to mouse, to assess the utility of GenomeMUSter to interpret and characterize human GWAS discoveries with mouse genetic traits. Hatoum et al. performed a multi-ancestry GWAS meta-analytic study to identify variants and genes broadly associated to use of multiple addictive substances humans and identified 42 significantly associated genes (Hatoum et al. 2023). Use of multiple substances simultaneously is the result of many aspects of response to environment, patterns of substance use, and effects of substance use. To interpret the role of human genetic variants in this process in light of the many hundreds of mouse traits that have been characterized in studies of phenomena associated with substance use disorders, we performed a meta-analysis of 397 mouse traits in the Mouse Phenome Database annotated to the Vertebrate Trait ontology term, VT:0010488, Response to Addictive Substance and separate meta-analyses of each of the child terms Response to Alcohol, Cocaine, Nicotine, and Withdrawal Response to Addictive Substance (VT:0010489, VT:0010718, VT:0010719, VT:0010721, and VT:0010722, respectively; **Fig. 5**). These phenotypic endpoints were obtained from 15 studies in the Mouse Phenome Database that measured the response to addictive substances. Substances studied included morphine, cocaine, methamphetamine, nicotine, and ethanol (alcohol) and measured various aspects of addiction-related endpoints, including sensitivity and tolerance, intravenous self-administration (IVSA) endpoints, observations related to balance, strength, cognition, and dexterity, bottle or lever choice, fear conditioning, as well as histopathology, and measures of lipids and metabolites (Chesler et al., Schoenrock et al. 2022, Bagley et al. 2022, Bubier et al. 2020, Wiltshire et al. 2015, Dickson et al. 2016, Crabbe et al. 1985, Crabbe et al. 2003, Rustay et al. 2003, Kamens et al. 2005, Metten et al. 2004, Metten and Crabbe 2005, Yoneyama et al. 2008, Thomsen and Caine 2011, Tsuchiya et al. 2012, Portugal et al. 2012, Kutlu et al. 2015). These cross-population analyses on the response to addictive substances were computed on ∼300 Diversity Outbred (J:DO) mice, Collaborative Cross, BXD recombinant inbred strains, and other classical and wild-derived inbred strains. Studies in these analyses that measured inbred strains included 6 to 45 strains (median of 8 strains).

**Fig 5.**
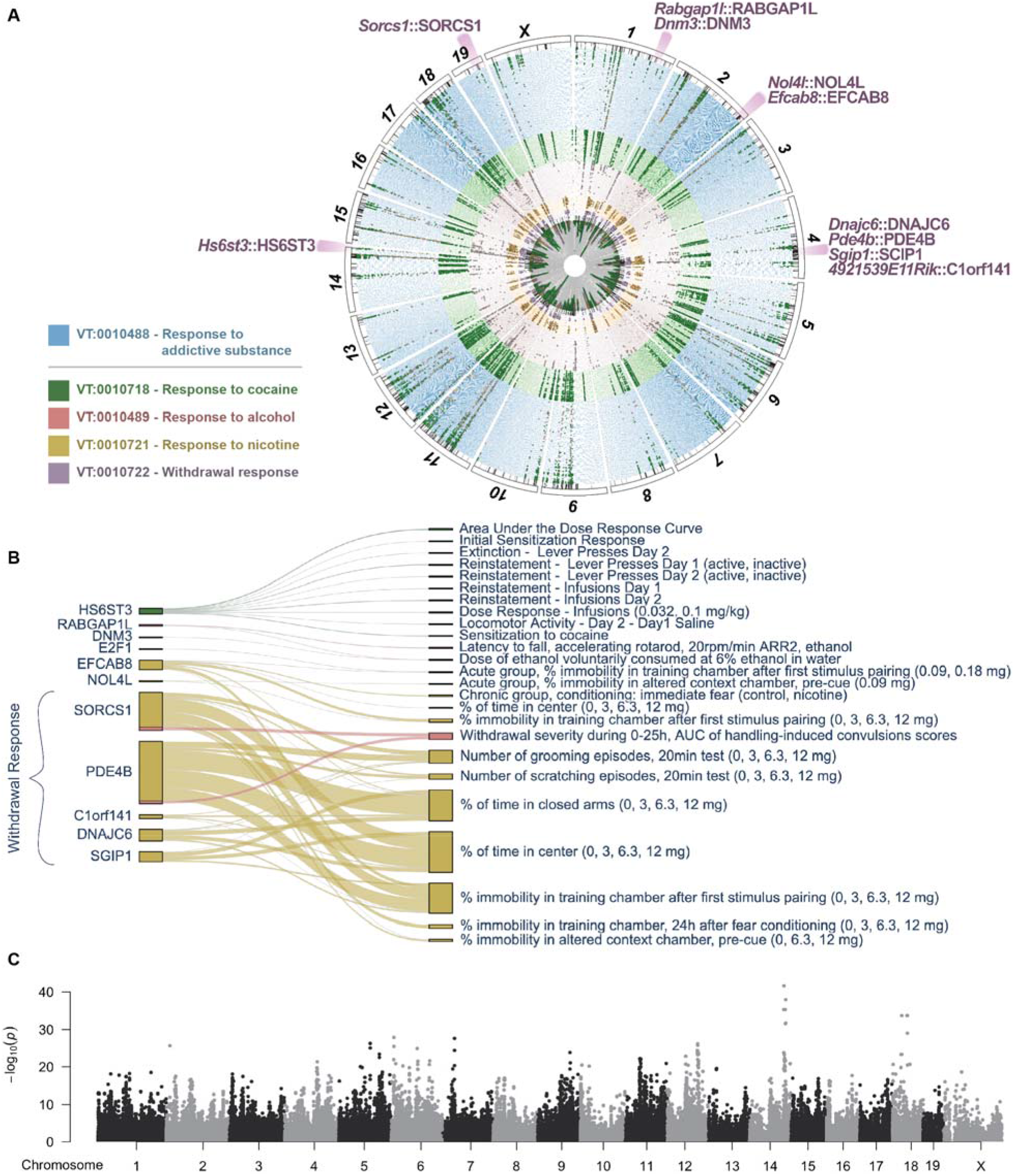
Cross-species integration of human multi-substance GWAS with mouse cross-population multi-trait meta-analyses of 390+ traits grouped by Vertebrate Trait (VT) ontology term. The circular plot (**A**) provides a bird’s eye view of these five mouse meta-analyses, variant effects, and associated pleiotropic genes detected in both mouse and human GWAS across multiple substances (Hatoum et al. 2023). Genomic regions of variants associated to these genes are highlighted, with associated mouse genes and their human orthologs annotated along the outer ring. The colors of each ring correspond to separate meta-analyses for Response to Addictive Substance (blue), Cocaine (green), Alcohol (red), Nicotine (orange), and Withdrawal Response to Addictive Substance (purple). Dots, colored by substance, indicate measures where the variant effect exists (m-value > 0.9; Han and Eskin 2011) to enable discernment between substance-specific effects and polysubstance effects. A full resolution version with annotations of all associated mouse genes is available (**Supplemental** Fig. 01) The Sankey plot (**B**) shows the number of variants affecting the mouse traits and the associated orthologous genes. The width of the line represents the number of variants and the color of the line represents the substance (green = cocaine, red = alcohol, orange = nicotine). The Manhattan plot (**C**) of the meta-analysis of the parent term, Response to Addictive Substance (VT: 0010488) on 397 mouse traits.

Across these five meta-analyses we detected a total of 5,705 variants with effects in the 99.5^th^ percentile or highly correlated with the variants in the 99.5^th^ percentile. We associated variants to mouse genes and their human orthologs as illustrated in Reynolds et al. These 5,705 variants were associated with 1,784 mouse genes and 1,850 of their human orthologs identified through additional data resources that include coding variation and eQTLs from GTEx (GTEx Consortium 2013), mouse striatum (Philip et al. 2023) and hippocampus (Skelly et al. 2019). Of these 5,705 variants, 1153 were detected in the meta-analysis of the parent term, Response to Addictive Substance (VT:0010488), and are associated with 525 human ortholog genes. Across the substance-specific meta-analytical results, 99 variants, associated with 65 human ortholog genes, were shared among multiple substances, indicating a possible role in general addiction processes. 51 of these 99 variants, associated with 41 human ortholog genes, were also detected both in the meta-analysis of the parent term, Response to Addictive Substance and in the meta-analysis of Withdrawal Response to Addictive Substance, indicating that these variants have a possible role in regulating a general mechanism of withdrawal across substances (**Fig. 5, Supplemental Fig. S1**).

These gene products were compared to two sets of human orthologue genes that were found to be significantly associated to multiple substance use disorders (Hatoum et al. 2023). Specifically, we used MAGMA (de Leeuw et al. 2015) to conduct whole-gene burden tests using summary statistics and also considered ‘mapped genes’ that were within 10kb of a genome-wide significant SNP as statistically significant. When compared with the results from Hatoum et al., our multi-substance meta-analyses in the mouse revealed HS6ST3, NOL4L, and PDE4B from the MAGMA-identified gene set and TNN, RABGAP1L, DNM3, EFCAB8, BPIFA2, E2F1, DNAJC6, PDE4B, SGIP1, C1orf141, and SORCS1 from the mapped gene set. One of the top hits, PDE4B is thought to affect the dopaminergic pathway (Snyder and Vanover 2017) and is one of the 15 genes that is associated with SNP effects observed across two human populations (Hatoum et al. 2023). By viewing meta-content for each of the mouse measures associated with *Pde4b*-related variation, we found that these were traits related to withdrawal from addictive substances and notably, the effect increased as the dose of the substance increased.

## DISCUSSION

Here, we present GenomeMUSter, intended to be the most comprehensive and interoperable, mouse genetic variation data resource service to date, containing more than 106.8 million genotyped, sequenced loci, or 70 billion imputed variant states for 657 inbred mouse strains representing common, wild-derived, recombinant inbred strains and their intercross derivatives. GenomeMUSter provides a comprehensive analytical SNP resource that can be used for a variety of research applications, which include characterizing the effect of genetic variants and their influence on transcripts and traits and refinement and resolution of genetic mapping experiments. It is important to note, however, that GenomeMUSter is not a static resource; rather, it is a dynamic and versioned resource that will continually be updated as new mouse sequence data is acquired. Because the sequenced and typed data used to build GenomeMUSter is on GRCm38 coordinates, we chose to keep GRCm38 coordinates for this version and provide access to the recently released Sanger (REL2021) dataset (Keene et al. 2011) in the GRCm39 assembly as well as an option to use liftOver to map GRCm38 coordinates onto GRCm39, while more rigorous annotation of the GRCm39 is in progress. Notably, assemblies will be continuously updated as more regions are sequenced and as we learn more of the structural variation present in the mouse with newer long-read sequencing-based assemblies. For example, long-read whole genome sequencing, as employed in the Stanford1 dataset (Arslan et al. 2023) included in GenomeMUSter and Ferraj et al. 2023, who sequenced a set of twenty diverse inbred strains from three sub-species of *Mus musculus*, provides rich information about structural variation in regions previously unexamined. Importantly, the analytical approach we present here can be applied not only to new mouse assemblies and new variation data but can also be applied to any other widely studied genetic model organisms, including *Drosophila*, rat, *C. elegans*, yeast, and many agricultural species for which incomplete reference genotypes are available (Mackay et al. 2012, Shimoyama et al. 2015, Sulston et al. 1992, Hillier et al. 2008, Güldener et al. 2005).

Imputation will ultimately be replaced by full sequence data for many strains but for legacy populations, heterogeneous stocks and inter-crosses, augmentation of sparse genotype data will still be required. Prior efforts provided high confidence imputations for only the well-characterized set of classical inbred strains, likely due to both the extreme sparseness of the data for some strains which were genotyped at as few as 1,638 loci and to the tremendous genetic diversity of the population, which limits the utility of known or inferred ancestry information due to a lack of similar cases that have sufficiently dense data in any given region. For example, while Wang et al. incorporated phylogenetic information, they excluded wild-derived strains because of the abundance of private alleles, which make it difficult or impossible to construct local haplotype blocks using the four-gamete rule in regions < 5kb. Moreover, some approaches that use HMMs require estimation of many parameters and importantly, the number of parameters to estimate in these methods increases linearly with the number of SNPs (Szatkiewicz et al. 2008). Given the lack of dense genotypes for the majority of strains and the number of private alleles among wild-derived strains, GenomeMUSter makes use of a larger genomic region on which to calculate phylogenetic similarity, does not require free parameter estimation, and employs the computationally efficient Viterbi algorithm, as implemented in HaploQA.

When examining the accuracy of the imputation algorithm, we found that for most strains it produced high imputation accuracies with a median imputation accuracy of 0.944 on the held-out test set. Lower imputation accuracies (< 0.70) in some wild-derived strains, such as PAHARI/EiJ, CAROLI/EiJ, and SPRET/EiJ (**Fig. 2**) were not unexpected as these strains are a different species (*M. pahari*, *M. caroli*, and *M. spretus*, respectively) from classical inbred strains (*M. musculus*) and the data for these strains was sparse in the datasets we included in GenomeMUSter. New full sequence data are being added, to improve the accuracy of information around the wild-derived strains (Thybert et al. 2018). The phylogenetic trees support a data-driven approach where imputed genotypes are based on known genotypes from phylogenetically similar predictor strains with limited co-occurring missingness and thus, if the most phylogenetically similar strains differ in important ways, i.e., are a different species, or the coverage in region is exceptionally sparse, we expect lower imputation accuracies. The phylogenetic trees also provide a rich genomic landscape to explore possible regions of cryptic genetic variation among strains. As more wild-derived inbred mouse strains are sequenced and included in GenomeMUSter, imputation accuracy for all strains will improve (Gambogi et al. 2023, Morgan 2022, Chang et al. 2017, Poltarak et al. 2018). A challenge with imputing genotypes for all missing allele states is that we inadvertently impute SNPs for insertions and deletions thus, known structural variation needs to be masked. Pangenome representations are a compact and even more scalable graphical means of presenting individual variation in the context of population diversity (Liao et al. 2023, Wang et al. 2022) that address this challenge, and will be utilized in future releases of GenomeMUSter.

GenomeMUSter serves as an analytical resource that is available programmatically for a wealth of applications in the study of genetic variation and phenotypic diversity in health and disease. For example, GenomeMUSter can be used for GWAS and genetic meta-analysis, to refine genetic loci underlying trait variation across mouse strain panels (Raza et al. 2023), and a host of other genome-scale operations. The extant inbred mouse strains have been broadly characterized over decades of mouse phenomic research. Examination of the large set of mouse traits that share orthologous regulatory variant targets provides a means to interpret and characterize GWAS variants.

We illustrated the utility of using GenomeMUSter for the genetic discovery of variants and orthologous genes for the complex diseases of addiction and Type 2 Diabetes. For example, our analysis of genes pleiotropic across human studies of addiction reveals that the GWAS candidate *Pde4b* most likely plays a role in withdrawal-mediated response to drug. It is speculated that this is indeed a broad mechanism of substance use disorder across all classes of drugs, as people often use substances to ameliorate the consequences of withdrawal, i.e. negative reinforcement is a general addiction process regardless of substance class. This is consistent with neurobiological mechanisms of substance use disorder (SUD) and provides evidence in support of the GWAS finding that PDE4B related variation is associated with addiction but not specific to any one class of substance. Notably, the interpretation provided through the use of multi-population, multi-trait meta-analyses in mice indicates future research directions in the functional characterization of human GWAS variants pharmacologically. Indeed, early clinical trials have already shown promise for *PDE4B* drug targets treating alcohol and opioid withdrawal (Burnette et al. 2021, Cooper et al. 2016).

We evaluated the use of GenomeMUSter in GWAS meta-analysis to discover genes and variants related to Type 2 Diabetes and obesity. We performed two analyses, the first using various physiological responses to a high-fat diet and the second, a meta-analysis of glucose-related endpoints which we predict would be more specific to Type 2 Diabetes. To test the veracity of these predictions we compared the gene to disease associations that were predicted in the metanalysis to curated annotations of human genes to Disease Ontology terms. We show that we were able to retrieve highly relevant terms using the human orthologs of mouse genes identified in our analyses. The most frequent Disease Ontology annotations of human orthologs of mouse genes associated with physiological responses to high-fat diet included Alzheimer’s Disease, followed by end stage renal disease, asthma, hepatocellular carcinoma, hypertension and obesity, and other inflammatory diseases. Each are known to be associated with Type 2 Diabetes (AD:Chatterjee and Mudher 2018, RD:Ritz et al. 1999, HC:Dyson et al. 2014, Hypertension:Okosun et al. 2001). The most frequent Disease Ontology annotations of human orthologs of mouse genes associated with variant effects on glucose related endpoints, a more specific diabetes related endophenotype, were Type 2 Diabetes, Alzheimer’s Disease and Obesity. This reveals that the multi-trait meta-analysis in mice was capable of identifying specific disease associated genes when a disease feature was used, and it was capable of detecting more general indirect environmental drivers of disease etiology such as high-fat diet, when this type of term was used. It should be noted that in meta-analysis, the quality of the result is driven by the study inclusion criteria. Broad search terms for disease related traits, while rapidly computationally reproducible and readily applicable to whole vocabularies and databases may result in false positive search results, and manual filtering, though labor intensive, may improve the precision of global associations of variants to disease. Use of specific mappings of traits to disease developed in efforts such as the Ontology of Biological Attributes (Stefancsik, et al. 2023) will further refine the precision of disease relevant queries of mouse trait data.

Human GWAS often connects variants to disease diagnoses using a very limited set of clinical assays. Mouse phenotyping studies combined with genomic harmonization through GenomeMUSter can have extensive breadth and depth, allowing an understanding of the specific processes and temporal involvement of genetic variants in the disease risk, manifestation, and progression. GenomeMUSter unlocks the utility of all genotyped inbred mouse populations for integrative, predictive genetic and genomic analyses by providing insight into previously unavailable genetic variation, enhancing the effective use of inbred mouse strains and establishing their versatility as disease models. Parallel efforts to refine the alignment of disease and trait vocabularies across species and identify variants with orthologous targets will greatly expand the potential for analytic operations using integrated genomics and phenomics in the laboratory mouse. GenomeMUSter and its accompanying analytical approach increase the scope, scale, and power to find relations among genetic variants and individual phenotypic variation and to use this variation to better model, map and characterize the variation associated with complex human disease.

## METHODS

### Pre-processing of whole genome sequence data

Whole genome sequence data of the Collaborative Cross (CC; Srivastava et al. 2017) were downloaded from the European Nucleotide Archive (ENA; https://www.ebi.ac.uk/ena/browser/view/PRJEB14673; Merged_69_flagged.tab.vcf.gz). Insertions and deletions (INDELS) and Multiple Nucleotide Polymorphism (MNP) alleles were excluded by filtering out loci with either INDEL or COMPLEX in the ‘INFO’ column. Single Nucleotide Polymorphism (SNP) alleles with read depths ≥ 5 were included. Heterozygous genotypes were called ‘H’ if both alleles met the allele depth requirement and accounted for at least 20% of the total read depth. This 20% threshold for heterozygous genotype percentage of total read depth was determined based on agreement with University of North Carolina (UNC) GMUGA genotyped SNP calls (Morgan et al. 2015) across chromosome 1.

We adopted the same filtering approach for read depth and calling heterozygous genotypes for the other whole genome sequencing variant sets, excluding INDELS and MNPs. For the C57BL/6J Eve (B6Eve; Sarsani et al. 2019) whole genome sequencing set, we included only genomic loci that had ‘PASS’ in the filter column. For the BXD recombinant inbred strains variant set, <u>we included the three lane C57BL/6J (∼80x)</u> and two-lane DBA/2J (∼100x) genotypes. The single lane genotype calls for the parental strains were not used. The long read whole genome sequencing variant set (Stanford1; Arslan et al. 2023) was processed similarly.

### Merging and harmonizing variant sets

GenomeMUSter harmonizes allelic state calls across 16 disparate datasets that include two Sanger releases (REL2004, REL1505, Keene et al. 2011), whole genome sequencing of the Collaborative Cross (CC) (Srivastava et al. 2017), BXD recombinant inbred strains (Sasani et al. 2022, Ashbrook et al. 2022), C57BL/6J Eve (B6Eve) (Sarsani et al. 2019), long read whole genome sequencing of 42 inbred strains (Arslan et al. 2023), and sparse genotypes from ten legacy datasets available from MPD (e.g., Kirby et al. 2010, Yang et al. 2011, Frazer et al. 2011, Morgan et al. 2015, Srivastava et al. 2017). Strain and allelic state data were merged by chromosome and within chromosome in 10 Mb segmented regions (**Fig. 5**). When there were disagreements among datasets regarding the genotype call for a strain and position, the consensus vote across all datasets was used. Genotypes were removed and imputed if no consensus was reached unless the disagreement was complementary, e.g., A and T.

### Imputation approach

Missing genotypes were imputed for each strain in each 10 Mb region spanning the genome using the Viterbi algorithm as implemented in HaploQA (https://haploqa.jax.org). Unlike other Hidden Markov Models (HMMs) where many free parameters must be estimated (Szatkiewicz et al. 2008, Kirby et al. 2010), the implementation of the Viterbi algorithm in HaploQA relies on given transition probability parameters, resulting in a computationally efficient estimate of the most probable path (**Table 2**). We incorporate local phylogenetic information without prior knowledge of relatedness in an approach that accounts for the degree of information in the region, i.e., co-occurring missingness. To impute genotypes on a target strain, we identified the optimal predictor strains, considered ‘haplotypes’ in the algorithm, by first selecting strains that had the fewest co-occurring missing genotypes with the target strain in the region, and second, had the smallest phylogenetic distance, and thus, were the most phylogenetically similar to the target strain in the region (**Fig. 5**). The local phylogenetic distance between each strain and all other strains was calculated per 10 Mb region using the merged variant set and the function ‘dist.dna’ in the R package ‘ape’ (Paradis and Schlep 2018) with model = ‘F84’ (Felsenstein and Churchill 1996). For each target strain in each 10 Mb region, potential predictor strains were ranked by the number of positions missing genotype calls that coincided with the positions missing genotype calls in the target strain. An alternative approach is to partition the genome based on local phylogeny (White et al. 2009) however, the sparsity of genotype-based analyses may yield limited benefits. We selected the twenty potential predictor strains with the most genotype information in the region that coincided with missing data in the target strain, i.e., had the lowest co-occurring missingness in the region. Next, we selected strains whose phylogenetic distance to the target strain in the region was lower than the 10^th^ percentile of phylogenetic distances between the target strain and all potential predictor strains. We required at least two predictor strains and thus, if no predictor strains could be chosen based on this threshold, we increased the phylogenetic distance threshold in increments of tenth percentiles until we found at least two predictor strains with the smallest overlapping missing genotypes that were phylogenetically closest to the target strain. While we required at least two predictor strains for each target strain, we aimed for at least four predictor strains for each target strain, including a maximum of twenty predictor strains.

### Imputation algorithm

For each target strain within a 10 Mb region, the major (A) and minor (B) alleles were determined across the optimal predictor strains for each locus. Using these classifications, genotypes for the target and predictor strains were translated to 1 for A if the genotype matched the major allele, 2 for B if the genotypes matched the minor allele, 3 if the genotype matched a third allele or H, and 0 for N if the genotype was unknown. The transition probability parameters in the HMM used by the Viterbi algorithm implemented in HaploQA are given, not estimated, and are based on expected genotyping or sequencing errors. Except for missing genotypes, the transition probabilities were set assuming there were minimal errors in the merged and harmonized variant set (**Supplemental Table S7**). (haplohmm.py in https://github.com/TheJacksonLaboratory/GenomeMUSter <u>derived from</u> https://github.com/TheJacksonLaboratory/haploqa).

### Validation of the imputation accuracy

To evaluate the accuracy of the imputation approach, a “hold out” test set for each strain was randomly selected in each 10 Mb region for 1000 locations or 10 percent (whichever was smaller) of known genotypes. This test set was not used in the imputation process. We calculated the accuracy on the held-out test set and set the confidence level in each 10 Mb region to the estimated accuracy on the held-out test set. If the imputation accuracy equaled 1, we set it to 0.999 to distinguish it from known genotype data. Because most strains had sparse data for the Y chromosome and mitochondria, we assessed overall accuracy for each strain over chromosomes 1-19, and X and provide a 95% Clopper-Pearson confidence interval. We calculated accuracy assuming that nucleotide complements were equivalent but provide both this and a traditional accuracy measure for completeness. After evaluating accuracy, allelic state data in the ‘hold out’ test sets were merged back into the full dataset. All analyses were completed in the R environment for statistical computing (R Core Team 2020), except the Viterbi algorithm in HaploQA.

### Validation of the utility of GenomeMUSter for multi-species integrative analyses

To evaluate the utility of this dense SNP resource for integrative genetic analysis, we performed multiple GWAS meta-analyses in two complex disease areas. First, we performed two cross-population meta-analyses on mouse traits that measured the physiological response to a high-fat diet and glucose related endpoints. Second, we performed five meta-analyses on addiction-related mouse traits annotated to Vertebrate Trait ontology terms in the Mouse Phenome Database. For all mouse meta-analyses, we first ran a GWAS mixed-effect model that accounts for population structure, i.e. kinship, (pyLMM; https://github.com/nickFurlotte/pylmm) on each distinct measure separately for animals of each sex. Next, we performed a mixed-effect meta-analysis (METASOFT; Han, Eskin 2011) on the summary statistics, as described in Bogue, Ball et al. 2023. We identified traits measuring the response to high-fat diet by searching ‘high-fat’ in the ‘intervention’ column of trait measures in the Mouse Phenome Database and similarly, for endpoints related to glucose, we searched the measure description column for ‘glucose’ (‘measurements.csv’ at https://phenome.jax.org/downloads). We identified addiction-related traits as those mouse traits annotated to Vertebrate Trait ontology term VT:0010488 Response to Addictive Substance and child terms Response to Alcohol, Cocaine, Nicotine, and Withdrawal Response to Addictive Substance (VT:0010489, VT:0010718, VT:0010719, VT:0010721, and VT:0010722, respectively). Prior to performing GWAS, we used the qtl2 R package (Broman et al. 2019) to impute genotypes of Diversity Outbred (DO) animals from the ‘Attie2’ study (Keller et al. 2018) and the ‘CSNA03’ study (Chesler et al.) onto a common SNP set so the GWAS summary statistics could be combined across inbred and outbred studies. (Bogue, Ball et al. 2023).

For each meta-analysis, we included variants in downstream analyses if they had effects in the 99.5^th^ percentile (top peaks) and then to account for linkage among variants, we also included variants near each peak that were strongly correlated with the peak locus. For each variant *i* and trait measure *j* in the meta-analysis, we calculated the *t*-statistic, 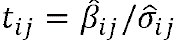 for each measure’s GWAS summary statistic: GWAS variant effect 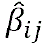, divided by the estimated standard deviation of the GWAS variant effect 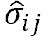, so for example, the meta-analysis on Response to Addictive Substance was computed on 484 GWAS summary statistics (estimated variant effects and standard errors) so each variant included in the meta-analysis had 484 *t*-statistics. A variant within 500,000 bps of a variant in the 99.5^th^ percentile (top peaks) was included if the Pearson’s *r* correlation coefficient between the variant’s *t*-statistics and the top peak variant’s *t*-statistics was ≥ 0.80. Variants detected in the meta-analyses were then passed through gene regulatory data resources, such as GTEx (GTEx Consortium, 2013), PLAC-Seq, ChIA-PET, and eQTL from Diversity Outbred mouse tissues, including striatum and hippocampus, to identify targeted mouse genes and their human orthologs (Reynolds et al. 2021).

We took two different approaches to assess the utility of using GenomeMUSter for GWAS meta-analyses for the discovery of variants and gene targets related to complex disease. For the meta-analyses on measures of the physiological response of exposure to a high-fat diet and glucose-related endpoints, we evaluated the human ortholog genes identified from these meta-analyses against the set of human diseases and conditions annotated to the human ortholog genes by the Alliance of Genome Resources (Kishore et al. 2020; Disease *Homo sapiens* associations at https://www.alliancegenome.org/downloads). For the addiction-related meta-analyses, we took the opposite approach, going from human to mouse. We compared the set of human genes associated with poly-substance use disorder through a cross-ancestry meta-analysis of observations from 1,025,550 individuals of European ancestry and 92,630 individuals of African ancestry (Hatoum et al. 2023) to the human ortholog genes identified through five meta-analyses of mouse measures annotated to Vertebrate Trait ontology terms in the Mouse Phenome Database. Two methods were used to assign significant genes from the human GWAS. First, in the European sample, we used MAGMA gene-based testing at the level of whole genes (de Leeuw et al. 2015), controlling for linage disequilibrium across genes and gene-size. Second, we took genome-wide SNPs that were significant in the European GWAS and the cross-ancestry GWAS and mapped them to any gene within 10kb of the significant gene for European, and directly on the gene for cross-ancestry (Hatoum et al. 2023).

### Accessing and using GenomeMUSter

GenomeMUSter and all individual datasets that went into building GenomeMUSter (**Table 1**) are stored in a specially constructed 8.1T instance of Google BigQuery, with a database schema consistent with many human datasets in BigQuery, e.g., 1000 Genomes Project (Siva 2008), ClinVar (Landrum et al. 2016), gnomAD (Koch 2020). The database was designed to enable efficient query of ∼70 billion genotypes across these strains and to be easily joined with elements from other datasets in BigQuery. This database is integrated with the publicly accessible MPD user interface (UI) and application programming interfaces (APIs), providing the research community ease of access to these data. Variant annotations to authoritative nomenclature, coordinates, and disease attributions are provided by the Mouse Variant Registry (MVAR) (https://mvar.jax.org). MVAR is an annotation service that evaluates submitted variant data to determine if any of the variants are equivalent to ones already in a repository (i.e., canonicalization). MVAR determines the type and consequence of variants using Jannovar (Jäger, et al. 2014). Associations of variants to phenotypes using terms in the Mammalian Phenotype Ontology (MPO) (Smith and Eppig 2009) are provided through expertly curated annotations from the Mouse Genome Database (MGD) (Blake et al. 2021).

GenomeMUSter and each of the individual datasets are publicly available via UI and API at https://muster.jax.org. Search inputs include gene symbols, Reference SNP identifiers (rsIDs), coordinates and/or coordinate ranges, and flanking regions. These inputs are used to search GenomeMUSter or optionally, the individual variant sets (**Table 1**) across the selected strains or strain panels. All data in GenomeMUSter is in Genome Reference Consortium Mouse Build 38 (GRCm38) coordinates however, we also provide access to Sanger SNP data in GRCm39 (REL2021) as well as an optional liftOver from GRCm38 to GRCm39 coordinates via pyliftover v0.4 (https://pypi.org/project/pyliftover/). Result tables (**Fig. 6A)** contain chromosome, location (bp), observed alleles, functional annotations from MVAR, gene symbol (MVAR), reference strain genotype data (classical C57BL/6J) followed by the remaining strains (default in alphabetical order). The confidence level for known genotypes is one and between 0 and 0.999 for imputed genotypes, and can be used to filter reports from the database. The confidence level represents the strain-specific accuracy on the held-out test set in each 10 Mb region. The confidence threshold is set to 1 only for observed genotype or sequence data. The confidence level can also be displayed as a heat map (**Fig. 6B)**. In addition to the simple download for small tables, larger queries are available via the API. GenomeMUSter can be accessed directly at https://muster.jax.org and via variant association tools in MPD (https://phenome.jax.org Bogue, Ball et al. 2023).

**Fig. 6.**
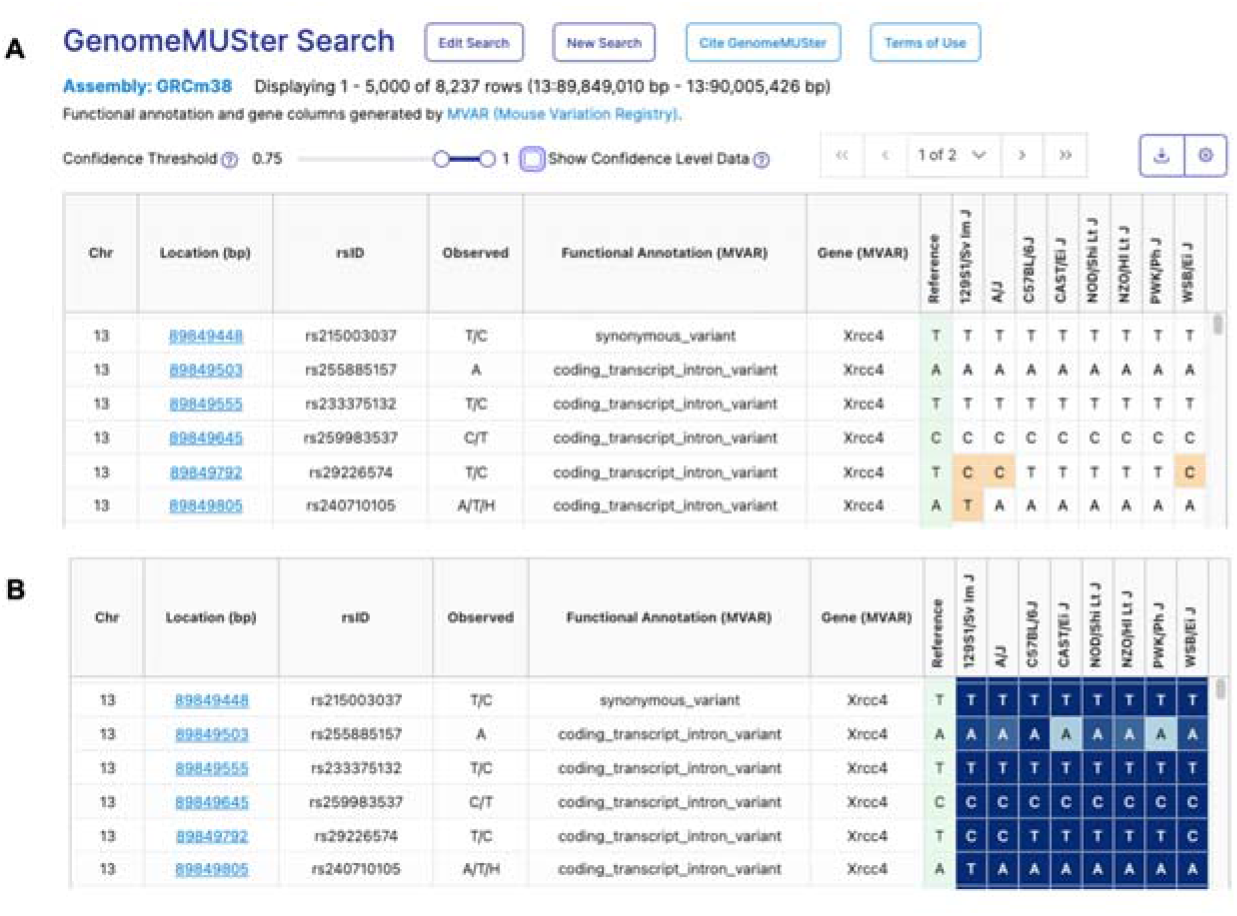
Example of GenomeMUSter search results. Users may choose not to show imputed genotype confidence levels (A) or to visualize the confidence levels of imputed genotypes (B) and filter results based on a confidence level threshold. In (B), lower confidence levels represented by lighter blue and higher confidence levels represented by darker blue.

GenomeMUSter is a dynamic and versioned resource, updated as new datasets become available. Additional genotype or sequence data submissions may be made to this public data resource by contacting.

## SUPPLEMENTAL MATERIAL

**Supplemental Table S1.** Coverage of GenomeMUSter and the 16 individual datasets per chromosome.

**Supplemental Table S2.** Coverage of GenomeMUSter and the 16 individual datasets across the genome.

**Supplemental Table S3.** Strain statistics aggregated across chromosomes 1-19 and X. Completeness and accuracy with 95% Clopper-Pearson confidence intervals.

**Supplemental Table S4.** Strain statistics for each chromosome. Completeness and accuracy with 95% Clopper-Pearson confidence intervals.

**Supplemental Table S5.** Disease Ontology terms annotated to human gene orthologs from a cross-population multi-trait GWAS meta-analysis of physiological response to a high-fat diet.

**Supplemental Table S6.** Disease Ontology terms annotated to human gene orthologs from a cross-population multi-trait GWAS meta-analysis of glucose related endpoints.

**Supplemental Table S7**. Transition probability parameters used by the Viterbi algorithm implemented in HaploQA for the genotype imputation approach. Code provided in haplohmm.py available at https://github.com/TheJacksonLaboratory/GenomeMUSter.

**Supplemental Fig. S1**. Mouse GWAS meta-analyses results of traits annotated to Vertebrate Trait ontology parent term VT:0010488 Response to Addictive Substance and child terms Response to Alcohol, Cocaine, Nicotine, and Withdrawal Response to Addictive Substance (VT:0010489, VT:0010718, VT:0010719, VT:0010721, and VT:0010722, respectively). The circular plot provides a bird’s eye view of these five meta-analyses, variant effects, and associated genes. Along the outer ring, mouse genes are annotated to each variant. Mouse genes in purple were also detected in multi-substance human GWAS (Hatoum et al. 2023). The colors of each ring and dots within rings correspond to separate meta-analyses for response to addictive substance (blue), cocaine (green), alcohol (red), nicotine (orange), and withdrawal response to addictive substance (purple). Dots indicate measures where the variant effect exists (m-value > 0.9; Han and Eskin 2011) to enable discernment between substance-specific effects and polysubstance effects.

## DATA ACCESS

The dataset generated for this paper and individual variant sets used to construct the merged and harmonized dataset are available at https://muster.jax.org. Code used to harmonize these data and impute missing allelic states is available at https://github.com/TheJacksonLaboratory/GenomeMUSter. Raw and processed whole genome sequencing data for Collaborative Cross (Srivastava et al. 2017) and the BXD dataset (Sasani et al. 2022) are available at the European Nucleotide Archive: PRJEB14673; Merged_69_flagged.tab.vcf.gz (CC) and PRJEB45429 (BXD). Raw whole genome sequencing data for B6Eve (Sarsani et al. 2019) and long read whole genome sequencing data for the Stanford1 dataset (Arslan et al. 2023) are available at the NCBI Short Read Archive database, PRJNA318985 (B6Eve) and PRJNA788143 (Stanford1).

## ACKNOWLEDGEMENTS

Funding provided by NIH DA028420 and by The Jackson Laboratory, The Cube Initiative Program Fund. We gratefully acknowledge Drs. Laura Reinholdt and Beth Dumont for editorial review of this manuscript and Dr. Reinholdt’s assistance with HaploQA. HaploQA was partially funded by the Mutant Mouse Resource and Research Center and Special Mouse Strain Resource at The Jackson Laboratory (U42 OD10921, P40 OD011102). Alexander S. Hatoum was supported by NIH K01 AA030083. David G. Ashbrook, Lu Lu, Robert W. Williams and the BXD sequencing effort were supported by the UT Center for Integrative and Translational Genomics and funds from the UT-ORNL Governor’s Chair. Gary Peltz and Fang Zhuoqing were supported by NIH/NIDA (1 U01 DA044399-01), awarded to Dr. Peltz. We also gratefully acknowledge the contributions of Lisa Tarantino, J. David Jentsch, Jane S. Adams, Stephen Grubb, Matthew Dunn, Benjamin L. Walton, Beena Kadakkuzha and members of the Computational Sciences Service at The Jackson Laboratory supported by the JAX Cancer Center Support Grant (P30 CA034196) for expert assistance with the work described in this publication.

## Author contributions

All authors reviewed the final manuscript. GAC, ZF, GP, LL, RWW, and DGA provided variation data and assisted in their incorporation into GenomeMUSter. AL, JB, MK, AKB, SD, BAS, and DOW contributed to the user interface design and development. BEK, FC, GK-R, and CJB provided the variant accessioning and annotations. RLB, HL, AS, MWG, HH, KS, JE performed the analysis and built the data platform. ASH and AA provided scientific expertise and data for the polysubstance use meta-analysis. RLB, MAB, VMP, EJC contributed the overall design and scientific leadership for this work.

## STATEMENTS AND DECLARATIONS

Nothing to declare.

## Notes

### Competing Interest Statement

The authors have declared no competing interest.

